# Emergence of increased frequency and severity of multiple infections by viruses due to spatial clustering of hosts

**DOI:** 10.1101/048876

**Authors:** Bradford P Taylor, Catherine J Penington, Joshua S. Weitz

## Abstract

Multiple virus particles can infect a target host cell. Such multiple infections (MIs) have significant and varied ecological and evolutionary consequences for both virus and host populations. Yet, the in situ rates and drivers of MIs in virus-microbe systems remain largely unknown. Here, we develop an individual-based model (IBM) of virus-microbe dynamics to probe how spatial interactions drive the frequency and nature of MIs. In our IBMs, we identify increasingly spatially correlated clusters of viruses given sufficient decreases viral movement. We also identify increasingly spatially correlated clusters of viruses and clusters of hosts given sufficient increases in viral infectivity. The emergence of clusters is associated with an increase in multiply infected hosts as compared to expectations from an analogous mean-field model. We also observe long-tails in the distribution of the multiplicity of infection (MOI) in contrast to mean-field expectations that such events are exponentially rare. We show that increases in both the frequency and severity of MIs occur when viruses invade a cluster of uninfected microbes. We contend that population-scale enhancement of MI arises from an aggregate of invasion dynamics over a distribution of microbe cluster sizes. Our work highlights the need to consider spatially explicit interactions as a potentially key driver underlying the ecology and evolution of virus-microbe communities.

## 1 Introduction

As some of the smallest and most abundant biological entities on earth, viruses lie at the foundation of many food webs [1]. Viruses drive biogeochemical cycles by turning over an estimated 20- 50% of all bacteria daily [2]. However, the interactions of individual viruses and microbial host cells are not well characterized *in situ* despite the magnitude of their aggregate effects. Only recently, the first signal of multiple infection (MI) of a host cell by viruses has been observed in natural settings [3]. This work used metagenomic analysis to identify viral genome signals from single-cell amplified genomes of microbes from oxygen-minimum marine zones. A subsequent analysis showed that MI is likely common with half of all sequenced bacterial genomes had evidence of MI [4]. A majority of coinfecting viruses in [4] derived from the same viral order, caudovirales. We interpret this signal as indicating high potential levels of MI by viral particles from related strains [5]. Nonetheless, it remains unknown what ecological factors drive rates of MI.

This lack of knowledge contrasts to a breadth of experimental work identifying examples of ecological and evolutionary consequences of MI. On the ecological side, life-history traits of viral infections such as the burst size [6, 7] and the latency time [8] can depend on the number of infecting viruses or multiplicity of infection (MOI). In fact, the fate of an infected cell can be MOI dependent when lysogeny is possible [9, 10]. On the evolutionary side, intracellular competition between multiply infecting viruses leads to a prisoner’s dilemma scenario in game theory [11]. “Defector” viruses utilize disproportionately large amount of intracellular resources compared to “cooperating” viruses. Defectors invade populations of cooperators inadvertently resulting in a reduction of the population-wide average fitness and possibly even viral population collapse [12, 13]. In extreme cases, exploitation of resource sharing leads to the emergence of viruses that completely rely on MI to propagate such as defective interfering particles, satellite viruses, and virophage [14, 15, 16]. These experimental studies conducted in shaken flasks, i.e., a well-mixed regime, do not provide insights as to mechanisms governing the rates of MI in complex environments.

Spatial epidemiological models have considered MI without an explicit link between cell death and viral release [17, 18]. In contrast, proposed models of viral dynamics with MI on individuals cells have focused in an immunological framework where viruses infect individual cells of a larger organism, without inclusion of explicit spatial effects [19, 20]. Prior spatial models of microbe-virus dynamics have considered plaque growth using PDEs [21, 22, 23, 24] and IBMs [25, 26] and the evolution of viral parameters using individual-based models (IBMs) [27, 28, 29, 30]. Only [27] included MI; however, the analysis did not quantify levels of MI and instead addressed whether MIs enhance virus-microbe coexistence. The question remains: how does realistic spatial clustering of populations alter subsequent MI dynamics?

Here, we address the basis for the emergence of MI using a stochastic, spatial IBM. We quantify the frequency of MI by comparing abundances of multiply infected hosts to abundances of singly infected hosts and abundances of viruses. Additionally, we characterize the severity of MIs by tracking the distribution of MOI across hosts. We then compare levels of MI between spatial and non-spatial models across parameter ranges that vary in spatial clustering. We find that MI frequency always increases with spatial clustering, whereas single infection frequency increases or decreases depending on which populations cluster. Similarly, MI severity increases with spatial clustering as displayed by fatter tails of the MOI distribution. Finally, we show how MI is enhanced during viral invasion of host clusters. As we discuss, the inclusion of spatial dynamics gives rise to both more frequent and more severe MIs, consistent with recent genomic-based inferences of environmentally sequenced microbes [3, 4].

## 2 Methods

### 2.1 Spatial model

We develop a stochastic, spatial IBM of virus-microbe dynamics. The domain is a twodimensional, periodic square lattice where at most one host and any number of viruses can occupy a lattice point. Dynamics occur at fixed time steps given stochastic processes that include cell growth, cell death, infection, lysis, and virus decay (see Supplementary Materials). Figure 1 shows how multiple infections can occur during the simulation. In this example, a colocated virus infects hosts at an average adsorption rate *ϕ*. Multiple infections occur if another colocated virus infects the previously infected host cell before lysis. Note, in a single time step more than one virus can infect the same host cell. Infected cells lyse at a rate *λ* releasing a burst size *β* viruses into the lattice point. Infected cells act as a sink for viruses since *β* is assumed here to be independent of MOI.

**Figure 1:**
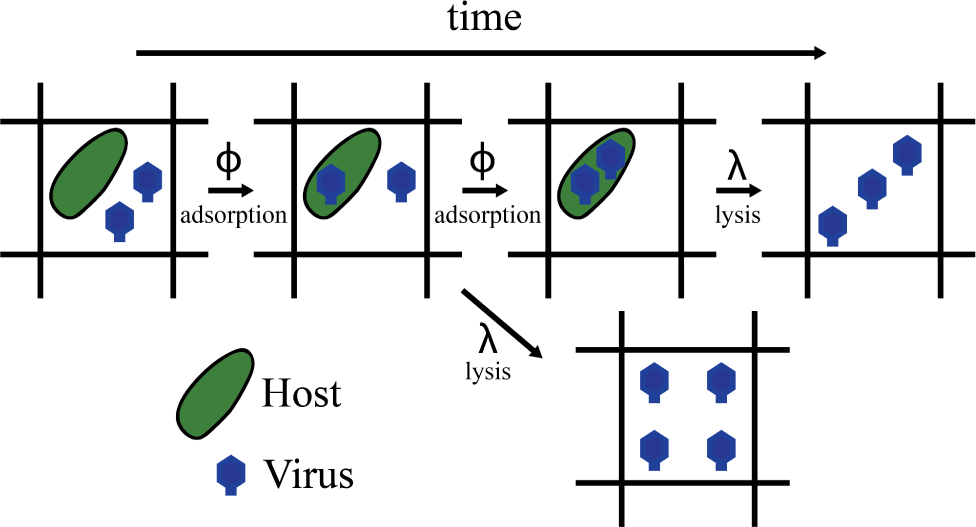
Infection dynamics in a single lattice point. Infection by a single virus occurs stochastically with adsorption rate, *ϕ*. The total rate of infection depends on the local viral abundance, *ϕV*_local_. Infected cells lyse with rate *λ* independent of MOI. Infected cells can be multiply infected if another infection event occurs before lysis. Lysis removes the host cell and replaces it with new viruses according to a fixed burst size independent of MOI. The burst size in this cartoon is 3 viruses for graphical convenience. We use a burst size of *β* = 20 in our models.

We initiate the spatial dynamics by randomly distributing hosts and viruses given that the total initial abundances match equilibrium solutions of the analogous mean field model. Each simulation is run for 105 timesteps corresponding to roughly 100 days given our simulation parameters. The goal was to simulate beyond transients such that our statistics would be representative of stationary distributions. The observed steady states are robust to initiating with alternative initial conditions.

### 2.2 Mean Field ODE model

Here we present a mean-field ODE model of virus-microbe dynamics with MI:

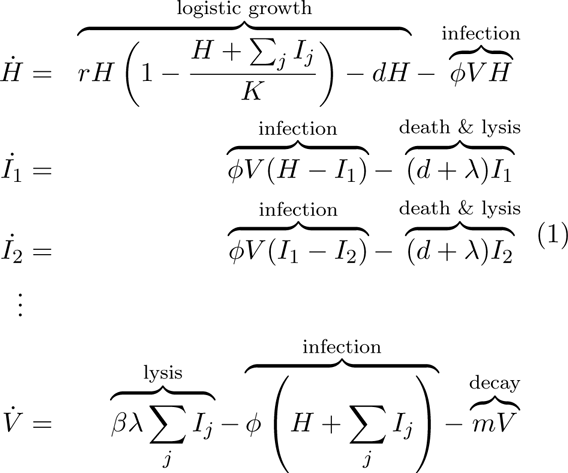

where *H* tracks uninfected hosts, *Ij* tracks infected hosts with MOI=j, and *V* tracks the viruses. We are interested in the dynamics of multiply infected cells, 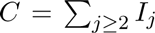 which follow:

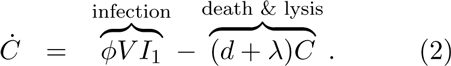

The parameters are the same between the spatial and mean field model. Differences between the two models result from spatial degrees of freedom and stochasticity. The steady-state solutions to the mean-field model can be solved exactly (see Supplementary Materials). The model assumes that both burst size and lysis rate are independent of MI.

We can obtain relevant MI statistics by solving the mean field model at equilibrium. From (2), we expect:

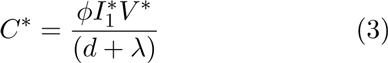

where * denotes abundances at steady-state. Similarly, solving 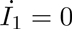 in (1) yields:

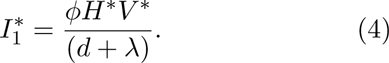

Solving (1) at equilibrium for any of the multiply infected host classes leads to a geometric sequence for the MOI distribution:

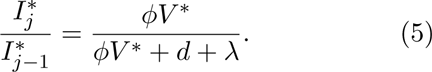

These derived results allow identifying deviations of the spatial model from the mean field expectation based on observed abundances.

## 3 Results

### 3.1 Spatial clustering emerges as viral adsorption and diffusion vary

We independently vary the adsorption rate *ϕ* and the viral diffusion constant *Dv* leading to emergent clustering in the spatial dynamics. The top two rows of Figure 2 show snapshots of the host and viral spatial dynamics respectively while only varying the adsorption rate, *ϕ*. Spatial clustering increases with increasing *ϕ* (columns moving left to right) as quantified by radial pair correlation profiles shown in the bottom row of Figure 2. These profiles display the ratio of the average number of hosts (viruses) located some distance away from each host (virus) over the expectation from random dispersal over the domain [31]. Values greater than 1 indicate increased frequency of hosts at a given distance. We define clustering to occur when the profiles are greater than a threshold value of 1.1 (dashed line). We chose this threshold as it is approximately the maximally observed 99% confidence interval when comparing to simulated ensembles of random dispersal. Low *ϕ* dynamics lead to cluster profiles indistinguishable from random dispersal of hosts and viruses (left column). As *ϕ* increases small clusters of hosts increase in frequency whereas the viral cluster profile is indistinguishable from random dispersal (middle column). High values of *ϕ*, where the dynamics are still robust against stochastic extinction events throughout our simulations, feature spatial clustering of both hosts and viruses (right column). Thus as *ϕ* varies we observe spatial dynamics feature no clustering, host clustering alone, then both host and viral clustering. We also varied the viral diffusion constant, *Dv*, which lead to increased viral clustering alone as *Dv* decreased (see Supplementary Materials). Interestingly, varying *Dv* did not lead to host clustering over the explored range. Viral clustering alone leads to temporary virus-free domains where hosts can locally reproduce to the local carrying capacity.

**Figure 2:**
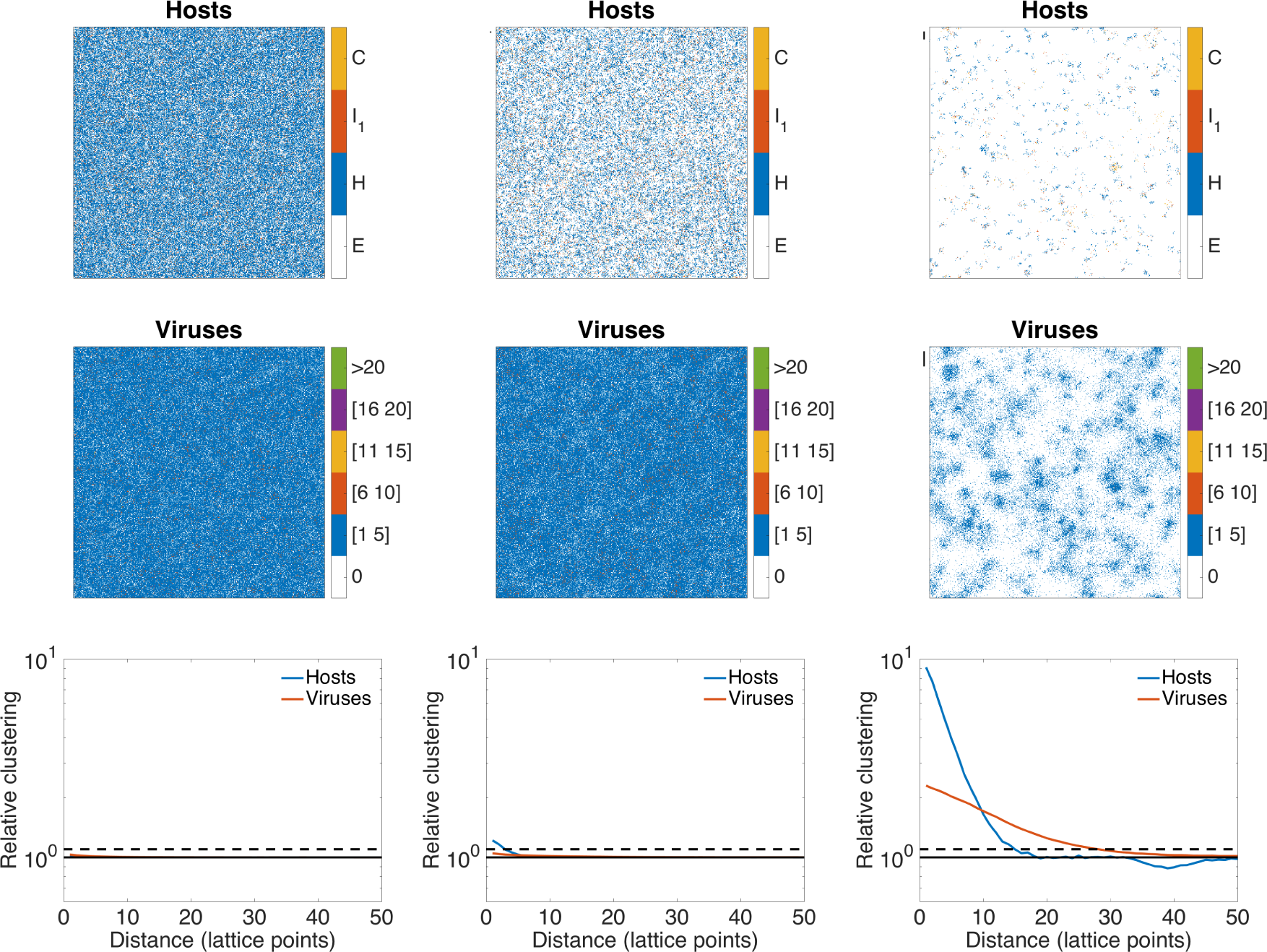
Spatial clustering increases with increasing adsorption. The color legend for hosts refers to multiply infected hosts (C), singly infected hosts (I1), uninfected hosts (H) and empty lattice sites (E). The color legend for viruses refers to the number of viruses located at each lattice point. (Left Column) Low adsorption, *ϕ* = 10^−8.4^ (ml/hr). (Middle Column) increased adsorption, *ϕ* = 10^−8.0^ (ml/hr) (Right Column) High stable value of adsorption, *ϕ* = 10^−7.1^ (ml/hr). Rows correspond to (Top) distribution of hosts, (Middle) distribution of viruses and (Bottom) radial pair correlation profile of hosts. The dotted line at *y* = 1.1 approximates the 99% confidence interval of the pair correlation profile when hosts and viruses are randomly dispersed. We use the intersection of this threshold line and the observed pair correlation profiles to define the cluster widths. When clustering occurs the corresponding cluster widths are plotted as black lines outside the top left corner of each of the spatial distribution plots. See supplementary material for other parameter values.

Independently increasing *ϕ* or decreasing *Dv* leads to increased clustering. The top row of Figure 3 shows this increase in clustering in terms of cluster widths. We define cluster widths as the maximal distance where the pair correlation profile exceeds the threshold. The range of parameters are colored according to whether the host, the virus, or both populations have nonzero cluster widths. Note, all data points from the simulations are averages over the last 10^4^ timesteps and over 10 replicates. The corresponding cluster widths for our prior examples are marked by black lines in the top two rows of Figure 2.

**Figure 3:**
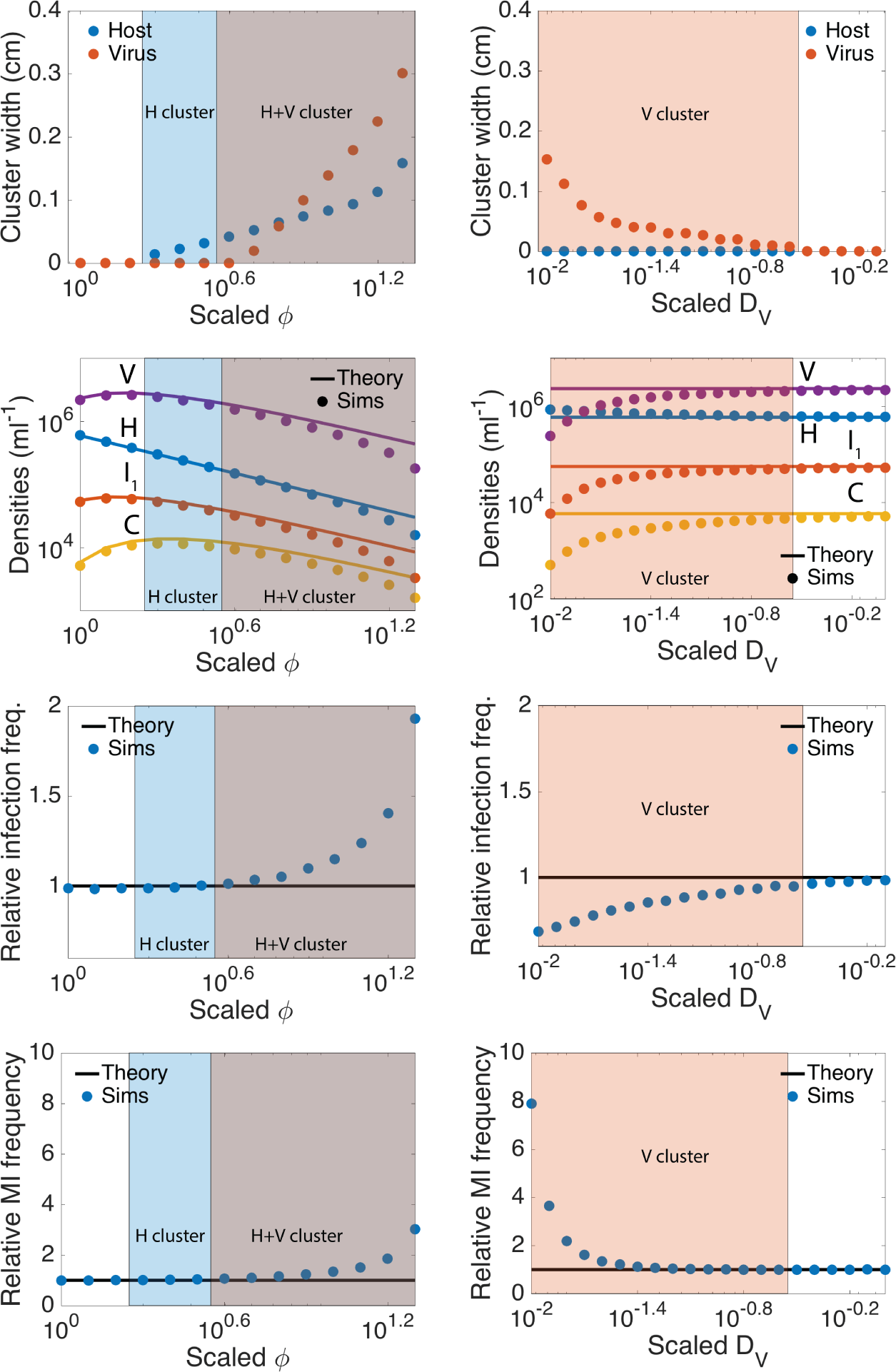
Emergence of clustering and its effects on densities and MI. The left column corresponds to results obtained by varying the adsorption rate and the right column corresponds to results obtained by varying the viral diffusion constant. The x-axes refer to scaling from reference parameter values of *ϕ* = 10^−8.4^ (ml/hr) and *Dv* = 2.04 × 10^−4^ (cm^2^/hr). All points are time and ensemble averages over the last 104 time steps and 10 replicate simulations. (First row) Cluster widths determined from non-zero x-value of the intersection between the pair correlation profiles and chosen threshold line. The transparent patches throughout indicate parameter values featuring clustering by hosts (blue), viruses (orange), or both (blue and orange overlay). (Second row) Population abundances where the line corresponds to solutions of the analogous non-spatial ODE model. (Third row) Relative frequency of single infections rate quantified by 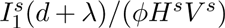 where the superscript “s” denotes abundance observed from the simulations. The dotted line is the mean-field expectation of unity. Points above (below) the dotted line indicate more (fewer) singly infected hosts given observed abundances of uninfected hosts and viruses. (Fourth row) Relative frequency of MIs rate quantified by 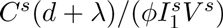 where the superscript “s” denotes abundance observed from the simulations. The dotted line is the mean-field expectation of unity. Points above (below) the dotted line indicate more (fewer) multiply infected hosts given observed abundances of singly infected hosts and viruses.

### 3.2 Effects of spatial clustering: increased relative rates of multiple infection

Here, we explore the effects of clustering on population densities and MI. To begin, we find that increased clustering leads to modest deviations in virus and host abundances from mean field expectations. The 2nd row of Figure 3 shows how most population abundances decrease from the mean-field expectation as a result of increased clustering. Only the uninfected host abundance increases compared to mean-field expectation as a result clustering when varying *Dv*. In that case, a smaller value in *Dv* limits viral movement leading to clustering of the viruses alone. Accordingly, the space between viral clusters acts as temporary domains where the hosts can grow uninhibited.

Spatial clustering leads to modest deviations in single infection statistics that depend on the form of clustering observed. The 3rd row of Figure 3 plots the relative infection frequency previously defined in the methods section and reiterated in the caption of Figure 3. Points above (below) the black line indicate larger (smaller) abundances of singly infected hosts as compared to the observed uninfected host abundances and the viral abundances. Increased clustering due to varying *ϕ* leads to increased relative infection frequencies. Meanwhile increased clustering due to varying *Dv* leads to decreased relative infection frequencies. This latter deviation is associated with negative spatial correlation between hosts and viruses due to increased fractions of the host population existing in temporary virus-free domains.

In contrast, clustering leads to significant increases in the MI frequency regardless of the form of clustering. The bottom row of Figure 3 shows the relative MI frequency previously defined in the methods section and reiterated in the caption of Figure 3. Points above the line indicate larger abundances of multiply infected hosts as compared to the observed singly infected host abundances and the viral abundances.

### 3.3 Effects of spatial clustering: MOI distributions with fatter tails

In this section, we show spatial clustering increases the severity of MIs as described by the MOI distribution – the abundance of hosts of increasing MOI. The MOI distribution is relevant when life-history traits can be MOI dependent (e.g., increased burst sizes, longer latency times). Our analysis gives a baseline understanding of the relevance of MOI skewing, though we do not explicitly modify life history traits as a function of MOI. The top rows of Figure 4 show the observed MOI distributions as compared to the previously derived mean-field expectation (black line) as we vary *ϕ*. We normalize the distributions by setting the density of singly infected hosts to 1. The observed MOI distributions match the mean-field geometric sequence for low *ϕ*. Spatial clustering leads to an increase in the mass of the tail of the MOI distribution, i.e., more hosts are infected by more viruses as compared to mean field. This deviation occurs continuously across parameter space as evidenced by the slight deviation present for weakly clustered dynamics (middle column Figure 4). The analogous plots when varying *Dv* are shown in the Supplementary Materials.

**Figure 4:**
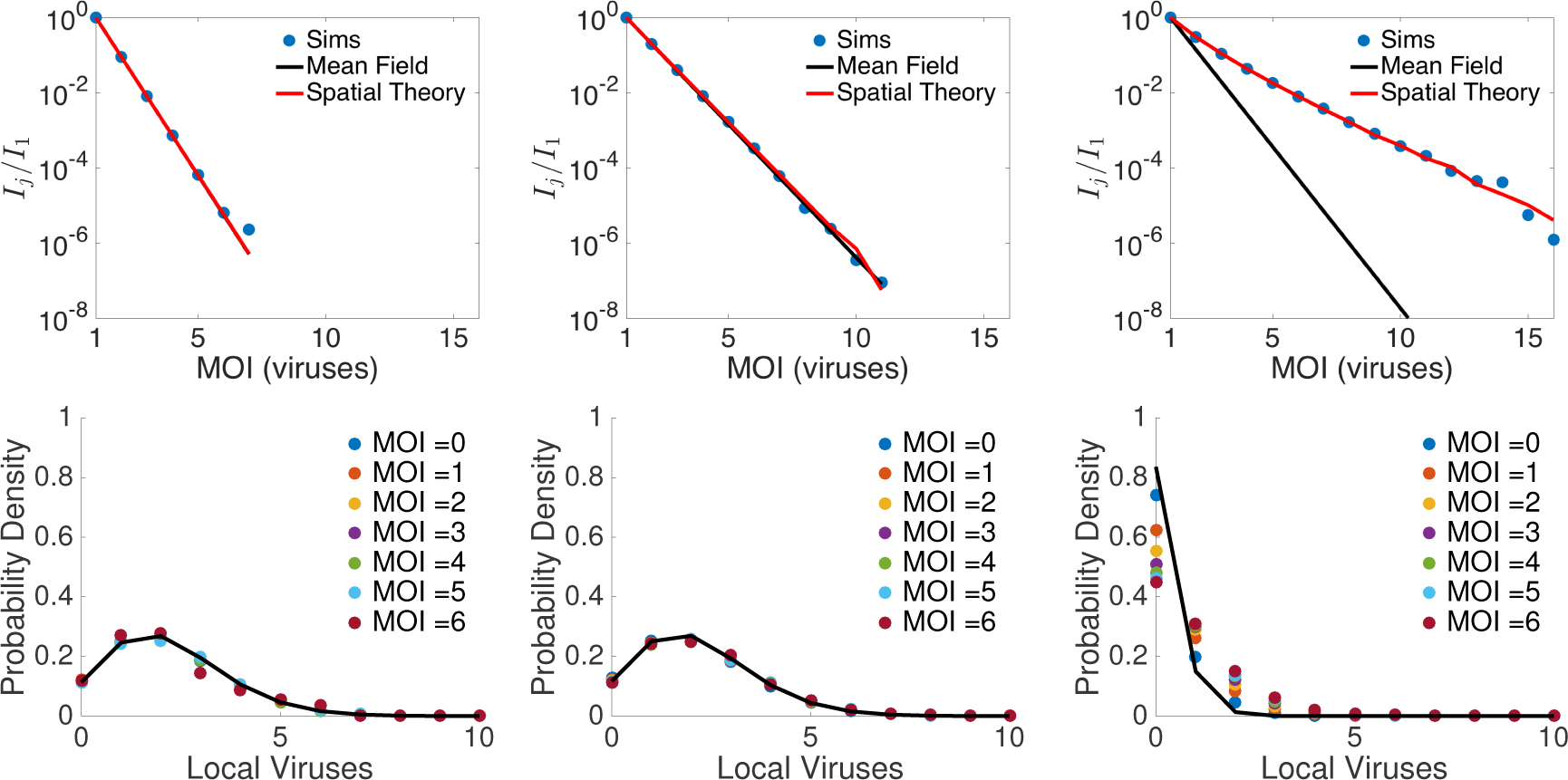
MOI distributions arising from local densities of viruses and underlying viral probability distributions (VPDs). VPDs quantify the local densities of viruses as normalized histograms of viral abundances colocated with host of a specified MOI. Predictions from spatial theory obtained using Eqn. 7. (Top) MOI distributions and (Bottom) normalized distributions of colocated viruses conditioned on MOI host for (Left Column) Low adsorption, *ϕ* = 10^−8.4^ (ml/hr), (Middle Column) increased adsorption, *ϕ* = 10^−8.0^ (ml/hr), and (Right Column) high stable value of adsorption, *ϕ* = 10^−7.1^ (ml/hr). Other parameters are the same as in Figure 2

The MOI distributions have relatively “fat” tails because more viruses are colocated with hosts of increasing MOI. The bottom row of Figure 4 shows the probability distributions of observing a number of external viruses in lattice points that contain hosts with a specific MOI. For clarity we only show these viral probability distributions (VPDs) up to MOI=6. Randomly distributing hosts and viruses across the domain leads to Poisson distributed VPDs (black lines) parametrized by the observed viral density. The observed VPDs match the Poisson distribution for all MOI in the low *ϕ*. Whereas for all other cases, the observed VPDs deviate from the Poisson distribution by skewing to the right, i.e., there are more viruses colocated with high MOI hosts than expected in the mean field theory. This skewing is more pronounced as clustering increases. The analogous plots when varying *Dv* are shown in the Supplementary Materials.

The viral distributions dictate the rate of flow between MOI types in the dynamics. The rate of viral infection in a lattice site linearly depends on the number of colocated external viruses. Here, we propose to adapt the mean-field approach, taking into account how local viral densities alter the dynamics. For example, the infection dynamics should follow

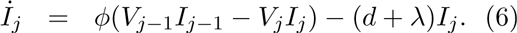

At equilibrium, this yields a sequence of multiplicative factors for the abundances of increasing MOI hosts:

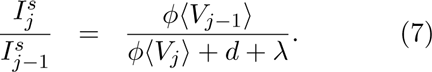

where the superscript “s” refers to the observed abundances from the spatial model that we conjecture to follow this relationship. We have replaced 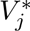 and 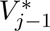 with 〈*V_j_*〉 and 〈*V*_*j–1*_〉 respectively, i.e., the means of the corresponding VPDs from the spatial IBM. The MOI distribution built from these scaling factors (red lines) matches the observed distributions in Figure 4.

### 3.4 Cluster invasion dynamics skew VPDs

We now show that viruses increasingly colocate with high MOI infected hosts during the viral invasion dynamics of a cluster of initially uninfected host cells. These invasion events become relatively more frequent with increased clustering and, in turn, further skew the statistics of the full system. For example, the top two rows of Figure 5 show a series of snapshot of the spatial dynamics with an adsorption rate 10^0.2^ ≈ 1.6 times the maximum of the range explored in Figure 3 (see Supplementary Materials for a corresponding movie). The typical dynamics at this parameter value oscillate wildly ultimately leading to stochastic extinction of at least one population within our simulation period. Here, most of the hosts and viruses are grouped in dense compact clusters (left frame). A single virus diffuses into a cluster initiating infection (middle frame). Note the source of this virus is not from the larger, discernible viral cluster in the left column. Rather, this virus remains from a previous cluster invasion event that occurred nearer. After lysis of the single infection event a cascade of infections leads to a wave of infections across the cluster (right frame). A large viral cluster remains after clearing the host cluster. Survival of the virus population relies on diffusing to and infecting a nearby growing host cluster before decay.

**Figure 5:**
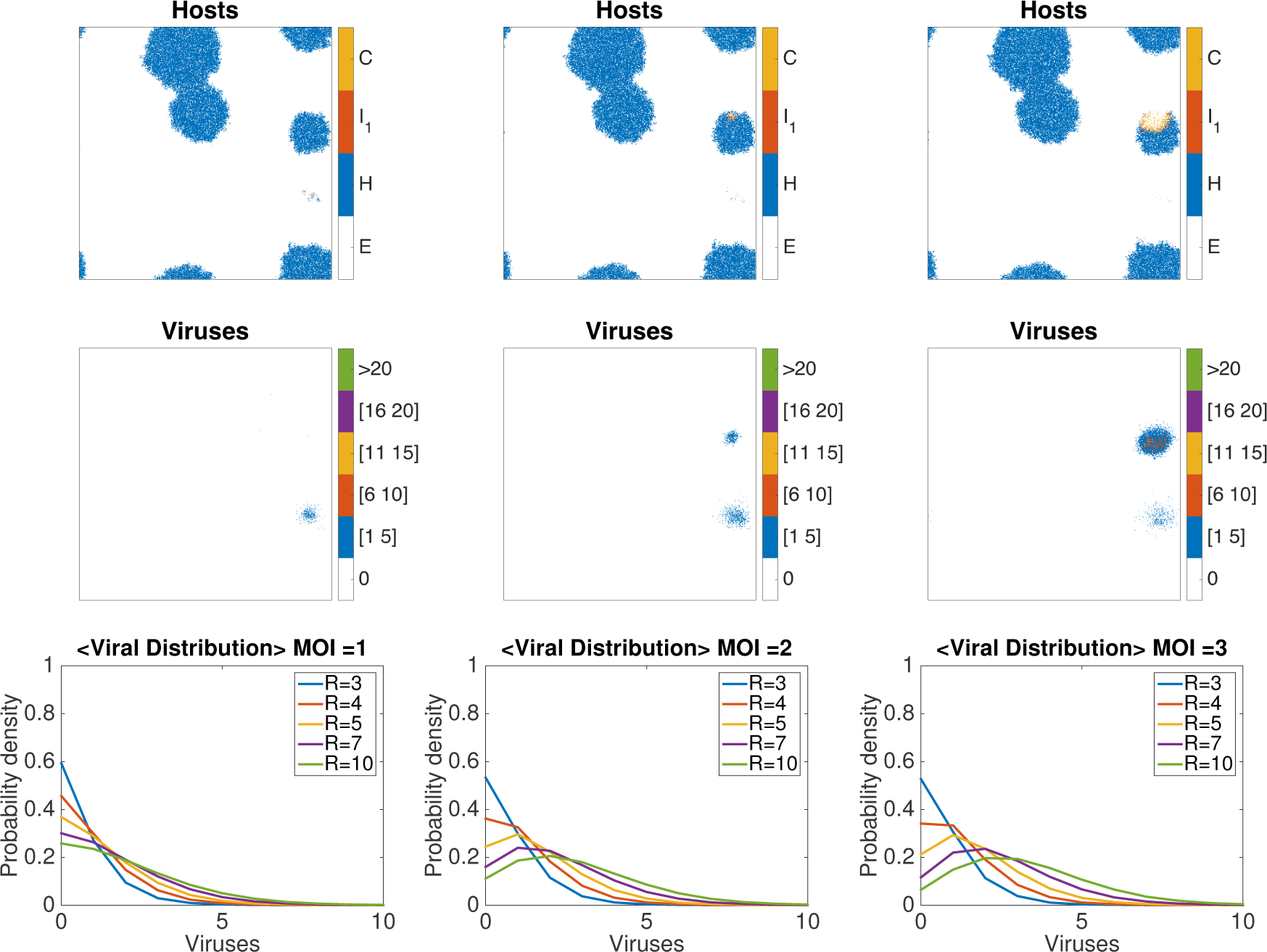
Virus infections spread wave-like across host clusters, enhancing MI. Snapshots of (Top) host and (Middle) viral dynamics when adsorption is increased such that unstable oscillations arise due to clustering (*ϕ* = 10^−6.9^ ml/hr). The dynamics feature the invasion of a host cluster by viruses over 50 times steps. The color legend for hosts refers to: C-multiply Infected hosts, I_1_singly infected hosts, H-uninfected hosts, E-empty lattice points. The color legend for viruses refers to the number of viruses located at each lattice point. (Bottom) Time averaged viral distributions for invasions of disc-shaped clusters of increasing radius (R) in units of lattice points. Each figure is conditioned on a different host MOI. The viral adsorption rate is *ϕ* = 10^−7.1^ (ml/hr) for the invasion dynamics.

In this example, the statistics and dynamics of the full system are completely determined by the growth and subsequent invasion of these clusters. To explore the effect of these dynamics on our MI statistics we simulate invasions on single host clusters. We consider maximally dense clusters of hosts of increasing radii and initiate a lytic event at the center. We utilize invasion simulations with immobile hosts that do not reproduce or diffuse to examine solely the effect of lysis and viral diffusion on MI. Figure 5 shows the time-averaged dynamics of the VPD for invasions of clusters with increasing radii. Each figure shows the VPDs of a specific MOI. For all MOI, the VPD skew to the right as the radius of the cluster increases, consistent with the results from the full stochastic IBM.

## 4 Discussion

In this paper, we demonstrated that both the frequency and severity of multiple infections (MIs) of microbial hosts by viruses increases due to spatial clustering. We identified the increase of MI frequency by comparing observed abundances of multiply infected host cells to abundances of singly infect host cells and viruses. The increased severity of MI was characterized by fatter tails in the multiplicity of infection (MOI) distribution. This fatter tail arises from positive skewing in the distribution of external viruses colocated with hosts of higher MOI types. Finally, we argued that invasion of larger host clusters leads to the skewed VPDs.

Part of the motivation for studying MI rates is that MIs can alter the dynamics of individual infections and, in turn, the entire population. While our models do not include MOI dependent parameters, our results do provide a baseline to compare future models where these feedbacks are included. For example, intracellular competition amongst multiply infecting viruses is more likely given spatially clustered dynamics. Such an increase in intracellular competition may lead to evolutionary conflict between viruses, e.g., a prisoner’s dilemma [11] and even to the extinction of a population [32]. In addition, the emergence of MI may also indicate when mechanisms to prevent secondary infections, like superinfection exclusion, should evolve [33]. Similarly, our model may also have implications for understanding the long-standing puzzle of persistence of multipartite viruses that require a high MOI for successful propagation [34].

While the patterns formed in our system are due to localized growth and limited dispersal, high density regions can occur by other means. The observed deviations between models arise from spatial correlations between host and viral types and are thus robust to specifics of how clustering forms. Hence, MIs could play a major role in shaping the population dynamics across a wide-range of patchy communities (e.g., in biofilms [35], during ocean blooms [36], resulting from chemotaxis [37] and resulting from turbulence [38], and standing patchiness [39, 40]). Furthermore, our observation of viral clustering in the absence of host clustering suggests that MIs may play a major role even in environments without observed microbial patchiness. Our simulations suggest this is more likely to occur for environments with a high host-viral ratio. Overall, observing rates of MI is of major empirical importance for understanding virus-microbe dynamics *in situ*. While recent work has demonstrated the existence of MIs in a targeted marine microbe [3] and within sequenced genomes [4] quantitative measurements of the frequencies are lacking.

In summary, this paper sheds light on how multiple infections emerge from population-scale dynamics taking place in spatially explicit domains. MOI dependent life-history traits can then act to modify subsequent population dynamics. It remains a question as to whether or not these kinds of feedback amplify or reduce MIs. In particular, MI allows for direct competition of viruses via shared resources inside the host. This complicates viral evolution as exploiting the host must be counterbalanced by exploitation from further viruses. Thus the effect of MI on viral evolution at a population level is relatively unexplored. The ubiquity of spatial clustering in natural environments suggests that increased attention on MI is necessary to understand the eco-evolutionary dynamics of the microbes and their viruses.

## Acknowledgements

This work was supported by NSF No. OCE1233760. We would like to thank Susanna Manrubia and Yu-Hui Lin for thoughtful comments during the preparation of the manuscript.

References

